# Natural self-attenuation of pathogenic viruses by deleting the silencing suppressor coding sequence for long-term plant-virus coexistence

**DOI:** 10.1101/2025.02.28.640734

**Authors:** Li Qin, Xiaoqing Wang, Zhaoji Dai, Wentao Shen, Fangfang Li, Aiming Wang, Adrián A. Valli, Hongguang Cui

**Author notes:** Correspondence (AAV); (HC). These authors contributed equally to this work (Li Qin, Xiaoqing Wang).

## Abstract

*Potyviridae* is the largest family of plant-infecting RNA viruses. All members of the family (potyvirids) have positive-sense single-stranded RNA genomes, with polyprotein processing as the main expression strategy. The 5’-proximal regions of their genomes encode two types of leader proteases: the serine protease P1 and the cysteine protease HCPro. However, their arrangement and sequence composition vary greatly among genera or even species. The leader proteases play multiple important roles in different potyvirid-host combinations, including RNA silencing suppression and virus transmission. Here, we report that viruses in the genus *Arepavirus*, which encode two HCPro leader proteases in tandem (HCPro1-HCPro2), can naturally lose the coding sequences for these two proteins during infection. Notably, this loss is associated with a shift in foliage symptoms from severe necrosis to mild chlorosis or even asymptomatic infections. Further analysis revealed that the deleted region is flanked by two short repeated sequences in the parental isolates, suggesting that recombination during virus replication likely drives this genomic deletion. Reverse genetic approaches confirmed that the loss of leader proteases weakens RNA silencing suppression and other critical functions. A field survey of areca palm trees displaying varied symptom severity identified a transitional stage in which full-length viruses and deletion mutants coexist in the same tree. Based on these findings, we propose a scenario in which full-length isolates drive robust infections and facilitate plant-to-plant transmission, eventually giving rise to leader protease-less variants that mitigate excessive damage to host trees, allowing long-term coexistence with the perennial host. To our knowledge, this is the first report of potyvirid self-attenuation via coding sequence loss.

**Author summary:** Plant viruses typically persist throughout the lifespan of their host plants, employing multiple strategies to ensure long-term survival. This study reveals an unusual self-attenuation mechanism in which potyvirids, likely through recombination, discard a large genomic fragment containing the RNA silencing suppressor coding sequence. This deletion reduces viral pathogenicity, enabling a peaceful long-term virus-plant coexistence. Meanwhile, the attenuated viruses may function as natural vaccines, protecting plants from reinfection by the full-length pathogenic strains or related variants. These findings highlight a direct link between viral evolution and long-term coexistence, suggesting that such adaptations may promote mutual benefits. Understanding these dynamics could provide valuable insights into virus-host interactions and inspire new approaches to plant disease management.

## Introduction

Plant viruses are strictly obligate intracellular parasites that survive and evolve within living host plants. Due to various constraints, their genomes are typically small, with limited coding capacity. As a result, they largely rely on host resources for replication, including the translation machinery, endomembrane systems (which form viral replication microenvironments), energy sources (*e.g.,* ATP), and host factors essential for assembling viral replication and movement complexes (Wang, 2015; Mäkinen et al., 2023; Pollari et al., 2024). To ensure successful infections, plant viruses have evolved a variety of strategies to counteract host’s multifaceted defense responses, which include RNA silencing, cellular autophagy, and hormone signal pathways (Chen et al., 2023; Ge et al., 2024; Li and Wang, 2019; Li et al., 2024; Wu et al., 2024; Yang et al., 2020).

Most animal-infecting viruses can be cleared from hosts by innate immune systems. In contrast, plant viruses typically persist within infected hosts throughout the whole lifespan. Plant viral infections can range from latent or mild to severe, but viruses rarely kill their natural hosts. This suggests that plant viruses have evolved sophisticated strategies to balance their infections, ensuring their survival without causing excessive damage to their hosts. Some of these strategies seem to be controlled by viruses themselves, as illustrated here by a few examples. Plant positive-sense RNA (+RNA) viruses undergo bottleneck effects and superinfection exclusion, which selectively allow just a few genomes to replicate and control the viral load at the cellular level (Zhang et al., 2018; Qu et al., 2020; Perdoncini Carvalho et al., 2022; Zhang et al., 2017). The presence of defective RNAs (D-RNAs), produced during the infection of several plant RNA viruses, interferes with viral replication, limiting virus overaccumulation (Simon et al., 2004; Budzyńska et al., 2022). Some viral proteins work as true infection modulators. For instance, the self-cleaving P1 protein regulates the release and, consequently, the activity of the RNA silencing suppressor (RSS) HCPro in most members of the *Potyviridae* family. This regulation prevents excessive virus accumulation at the onset of infection and mitigates the overactivation of defense mechanisms, ultimately facilitating long-term infection (Pasin et al., 2014; Pasin et al., 2020). Moreover, viral-encoded proteases involved in the proper viral polyprotein maturation have also been shown to cleave host factors, thereby triggering plant immunity to limit viral accumulation (Jia et al., 2025). Interestingly, it has been demonstrated that the genome sequence of a plant virus can be manipulated using synonymous mutations to outcompete its wild-type counterpart (Gonzalez de Prádena et al., 2020), further supporting the idea that virus accumulation within plants is not necessarily maximized.

*Potyviridae* is the largest family of RNA viruses infecting plants, comprising 235 species described to date classified into 12 genera. This family includes several agriculturally relevant viruses, such as turnip mosaic virus (TuMV), plum pox virus (PPV), potato virus Y (PVY), and soybean mosaic virus (SMV), among others (Cui & Wang, 2019; Yang et al., 2021; Inoue-Nagata et al., 2022; Mäkinen et al., 2023; Pollari et al., 2024). Members of the *Potyviridae* family, known as potyvirids, have positive-sense single-stranded RNA (+ssRNA) genomes ranging from 8.2-11.5 kb, encapsidated within flexuous filamentous particles (11-20 nm × 650-950 nm). Although the majority of potyvirids are monopartite, members of the genus *Bymovirus* have two +ssRNA genomes (RNA1: 7.2-7.6 kb; RNA2: 2.3-3.7 kb), forming filamentous particles with the modal lengths (250-300 nm, 500-600 nm) (Inoue-Nagata et al., 2022). Monopartite genomes contain a large open reading frame (ORF) and another short ORF, termed *pipo* (Chung et al., 2008). During replication, viral RNA polymerase slips at very low frequency in a conserved motif (G_1-2_A_6-7_) in the middle of P3 coding sequence to generate a small fraction of frame-shifted genomic RNA (Olspert et al., 2015; Rodamilans et al., 2015; White, 2015), in which the *pipo* ORF is now in frame with its upstream sequence. Upon translation, a large polyprotein and a shorter one are generated and proteolytically processed by virus-encoded proteases (*e.g.*, P1, HCPro and NIa-Pro) into 10 to 12 mature functional factors (Cui & Wang, 2019). Remarkably, recent reports have increased our knowledge on the coding capacity of potyvirids: (ⅰ) the antisense genome may encode additional small peptides, as in the case of the rORF2 from TuMV, which was experimentally proven to be essential for viral infection (Gong et al., 2023; Li et al., 2024; Cheng et al., 2024); and (ⅱ) transcriptional slippage motifs are more flexible than originally thought, and they are widespread across potyvirid genomes, leading to the production of truncated proteins or novel proteins whose functions remain to be investigated (Valli et al., 2024). The central and 3’-terminal regions of monopartite potyvirid genomes, which correspond to the RNA1 of bymoviruses, are relatively conserved in gene arrangement and sequence composition. This region encodes essential viral factors: P3, P3N-PIPO, 6K1, CI, 6K2, VPg, NIa-Pro, NIb, and CP. Each of these proteins play multifunctional roles in viral replication, movement and encapsidation (Cui & Wang, 2019; Yang et al., 2021). The 5’-proximal regions of potyvirid genomes encodes leader proteases, which vary among genera. These include the serine protease P1 and the cysteine protease HCPro (Cui & Wang, 2019). Notably, RNA2 in bymoviruses encodes two proteins: P1 (a homolog of HCPro) and P2 (Valli et al., 2018; Pasin et al., 2022). The arrangement of leader proteases varies among different genera (even within the same genus), including configurations such as P1-HCPro, a single P1, P1a-P1b, a single HCPro, and HCPro1-HCPro2 (Pasin et al., 2022; Hu et al., 2023). To date, several functions have been assigned to these proteases (Valli et al., 2018; Cui & Wang, 2019), including: (i) polyprotein maturation, as both P1 and HCPro have *cis*-cleavage activity for their own release from virus-encoded polyproteins; (ii) RNA silencing suppression, given that at least one leader protease in each potyvirid works as RSS, with HCPro-like proteins playing this role in most cases, including the P1 in bymoviruses and HCPro2 in arepaviruses (Chen et al., 2023; Qin et al., 2021); (iii) host range determination and adaptation, which is in line with its huge sequence variability (Gou et al., 2023; Shan et al., 2015, 2018; Salvador et al., 2008; Valli et al., 2007); and (iv) vector-mediated transmission between plants, as HCPro proteins from all tested potyvirids facilitate vector transmission by presumably acting as a molecular bridge between viral particles and vectors (Govier & Kassanis, 1974; Stenger et al., 2005; Valli et al., 2018; Szydło et al., 2024).

*Arepavirus* is a newly established genus in the family *Potyviridae*, comprising two definitive viruses: *Areca palm necrotic spindle-spot virus* (ANSSV) and *Areca palm necrotic ringspot virus* (ANRSV) (Yang et al., 2018, 2019; Inoue-Nagata et al., 2022). Both viruses infect in nature areca palm (*Areca catechu L.*), a perennial palmaceous plant with high medical value, causing severe foliage chlorosis and necrosis. The ANRSV is highly epidemic in the main growing regions of Hainan, China, with an average incidence rate of 19% and many identified isolates; in contrast, only one ANSSV isolate (ANSSV-HNBT) has been identified in the field. Notably, both viruses share a unique leader protease arrangement, featuring two HC-Pro proteins in tandem (HCPro1-HCPro2) (Qin et al., 2021; Cui & Wang, 2019). Beyond its role in *cis*-cleavage and RNA silencing suppression (Qin et al., 2021), HCPro2 also facilitates virus movement between cells. This process is coordinated with two other viral factors (CI and CP) and the host Rubisco small subunit (RbCS) (Qin et al., 2024).

Since 2018, we have observed that some areca palm trees in the field gradually recovered from arepavirus-induced disease, with severe foliage necrosis becoming mild or even disappearing. This intriguing phenomenon caught our attention, prompting us to investigate the underlaying mechanisms of recovery. In this study, we demonstrate that recovered trees remain infected with arepaviruses, even though viruses harbor a significant genomic deletion. Further analysis reveals that these viruses have lost the coding sequence for their leader proteases, likely due to recombination events. Additionally, we also show that the absence of leader proteases weakens viral RNA silencing suppression, as well as other important functions. However, despite this attenuation, viruses retain the ability to replicate and move between cells. Based on these findings, we propose that viral genomic deletion serves as a natural self-attenuation mechanism, likely promoting long-term plant-virus coexistence with mutual benefits.

## Results

### Recovery of an areca palm tree infected with ANSSV is associated with a drop in viral load and a deletion of HCPro1-HCPro2 coding sequence in the viral genome

ANSSV is the first described and type species in the genus *Arepavirus* (Yang et al., 2018; Inoue-Nagata et al., 2022). The virus was first identified in 2017 in a diseased areca palm tree showing severe foliage necrosis and chlorosis symptoms in Baoting, Hainan, China (Fig 1A). The full-length genomic sequence of the ANSSV-HNBT isolate from 2017 (referred to as ANSSV-BT17 in this study) was determined by conventional cloning and sequencing (Accession number in GenBank database: MH330686). Intriguingly, the symptoms in newly developed leaves of that particular tree gradually faded over time. As shown in Fig 1A, by 2023, only mild chlorosis symptoms were observed in the upper leaves. This intriguing phenomenon attracted our attention and prompted us to investigate the underlying mechanism(s) of disease recovery.

**Fig 1.**
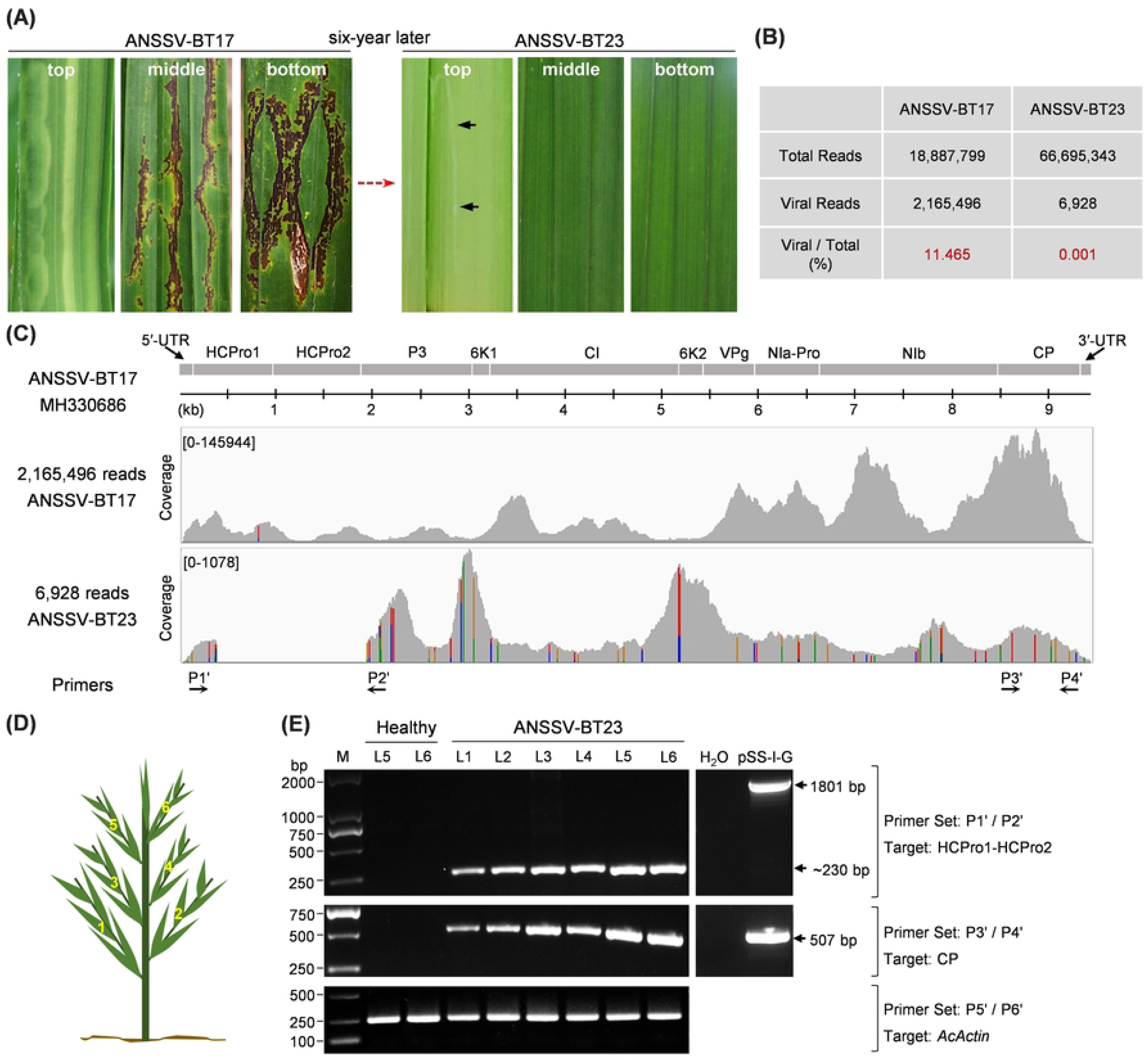
ANSSV lost nearly the complete coding region of HCPro1-HCPro2 during infection in an areca palm tree. (A) Leaf symptoms of an ANSSV-infected tree observed in 2017 and 2023. Virus isolates from the tree in 2017 and 2023 were designated as ANSSV-17 and ANSSV-23, respectively. Mild chlorosis is indicated by black arrows. (B) Summary of clean reads derived from RNA-seq data corresponding to the infected tree in 2017 and in 2023. (C) Alignment of clean reads with ANSSV-BT17 genome (MH330686). RNA-seq data from the infected tree in 2017 (upper panel) and in 2023 (lower panel) were used for the alignment. The horizontal axes represent nucleotide positions in the ANSSV-BT17 genome, while vertical axes indicate coverage. Nucleotide differences from the reference genome are highlighted with color bars: cytosine in blue, guanine in brown, adenine in green, and uridine in red. (D, E) RT-PCR analysis of RNA samples from the infected tree in 2023 using the indicated primer pairs. All leaves in the infected tree, L1-L6 (D), were tested. Equivalent amplicons from pSS-I-G, the ANSSV-BT17 infectious cDNA clone (Qin et al., 2021), were used as control. The RT-PCR product corresponding to *Areca catechu Actin* (*AcActin*) served as an internal control (lower panel).

Firstly, we performed RNA-seq to fully sequence ANSSV from the tree using samples harvested in 2017 (strong disease symptoms) and in 2023 (mild chlorosis). In the 2017 sample, 11.465% of the total reads aligned to the genomic sequence of ANSSV-BT17 (Fig 1B and 1C). However, in the 2023 sample, only 0.001 % of total reads mapped to the viral genome (Fig 1B and 1C), indicating a significant decrease in viral load over the time. We then compared the alignment maps of virus-derived reads from 2017 and 2023 against the ANSSV genome. Surprisingly, a large portion of the HCPro1-HCPro2-coding region was absent in reads from the sample harvested in 2023 (Fig 1C), suggesting that nearly the entire coding sequence of HCPro1-HCPro2 had been lost. We refer to this ANSSV variant carrying the genomic deletion as ANSSV-BT23. To confirm this finding, we designed two pairs of primers to perform reverse-transcription polymerase chain reaction (RT-PCR): the P1’/P2’ pair flanks the lost genomic region in ANSSV-BT23, whereas P3’/P4’ spans the CP coding region of this virus (Fig 1C). RNA was extracted from independent leaf samples collected in 2023 and used as a template for RT-PCR reactions (Fig 1D). The results showed that the CP region was amplified from RNA of all tested samples, confirming the presence of ANSSV throughout the plant (Fig 1E). As expected, an 1801-bp amplicon was produced when the infectious cDNA clone of ANSSV-BT17 (pSS-I-G) (Qin et al., 2021) was used as template. In contrast, only a short amplicon (∼230-bp) was consistently yielded from RNA of all leaf samples collected in 2023 (Fig 1E). Taken together, these results demonstrate that the entire tree is infected with ANSSV-BT23 variant, which has lost nearly the entire HCPro1-HCPro2 coding sequence.

#### The deletion of the HCPro1-HCPro2 coding sequence also occurs in the ANRSV genome

The fading of severe necrosis symptoms in newly developed leaves was also observed in ANRSV-infected trees in the field. In 2023, we selected an areca palm tree in Haikou, China, where symptoms had fully disappeared, to assess the presence or absence of HCPro1-HCPro2 in ANRSV (Fig 2A). This tree had showed severe foliage chlorosis and necrosis in 2021 (Fig 2A), however, we did not collect samples for virus sequencing at that time. Leaf samples were collected in 2023 and used to prepare RNA for RNA-seq analysis. The results showed that 0.03% of total reads mapped to the reference genome of ANRSV-XC1 (Accession no. MH395371), which had been fully sequenced by conventional methods (Yang et al., 2019). Surprisingly, no reads aligned against the 5′ region of ANRSV-XC1 genome (Fig 2B), suggesting that this ANRSV variant, here referred to as ANRSV-HK23, has suffered a significative deletion that includes the HCPro1-HCPro2 coding region.

**Fig 2.**
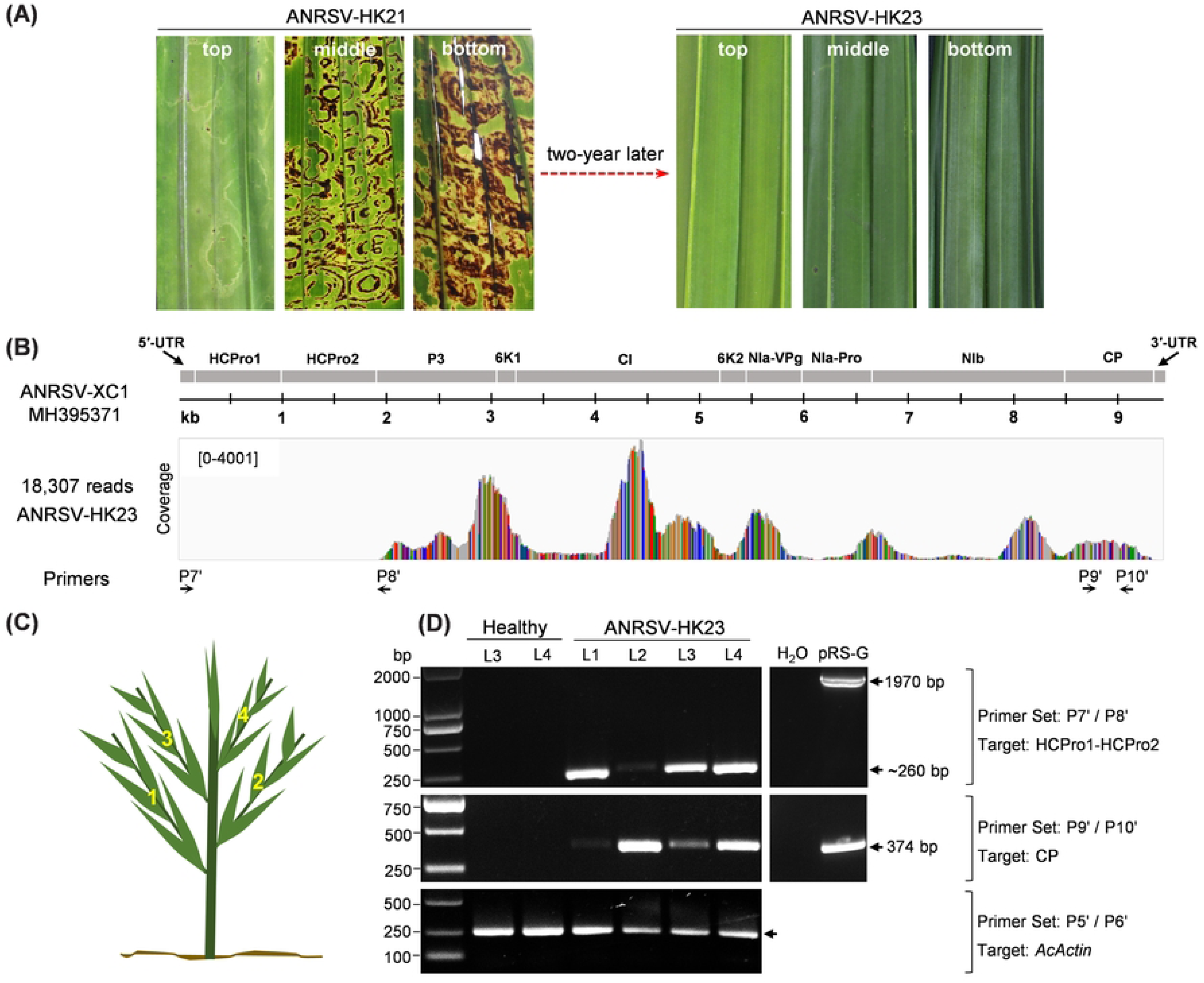
ANRSV lost the complete coding region of HCPro1-HCPro2 during infection in an areca palm tree. (A) Leaf symptoms of an ANRSV-infected tree observed in 2021 and 2023. Virus isolates from the tree in 2021 and 2023 were designated as ANRSV-HK21 and ANRSV-HK23, respectively. (B) Alignment of clean reads with ANRSV-XC1 genome (MH395371). RNA-seq data from the infected tree in 2023 were used for the alignment. The horizontal axes represent nucleotide positions in the ANRSV-XC1 genome, while vertical axes indicate coverage. Nucleotide differences from the reference genome are highlighted with color bars: cytosine in blue, guanine in brown, adenine in green, and uridine in red. (C, D) RT-PCR analysis of RNA samples from the infected tree in 2023 using the indicated primer pairs. All leaves in the infected tree, L1-L4 (C), were tested. Equivalent amplicons from pRS-G, the ANRSV-ZYZ infectious cDNA clone (Wang et al., 2021), were used as control. The RT-PCR product corresponding to *Areca catechu Actin* (*AcActin*) served as an internal control (lower panel).

Next, we followed a similar approach to the one described above to validate the RNA-seq results using RT-PCR. Therefore, we designed two pairs of primers based on ANRSV-HK23: P7’/P8’ pair flanks the genomic region corresponding to the potential deletion identified by RNA-seq (including the HCPro1-HCPro2 coding sequence), and P9’/P10’ pair targets the CP coding sequence of this virus (Fig 2B). All four leaves from the tree in which ANRSV-HK23 was identified were sampled for RNA extraction and further RT-PCR analysis (Fig 2C). Amplicons corresponding to a fragment of the CP coding region were successfully amplified using the P9’/P10’ primer pair (Fig 2D), confirming the presence of an ANRSV variant throughout the entire tree. As expected, when the infectious cDNA clone of ANRSV-ZYZ (pRS-G) (Wang et al., 2021) was used as template, a 1970-bp amplicon was produced. In contrast, a much shorter amplicon (∼ 260 bp) was generated with the P7’/P8’ primer set when using RNA samples from the infected tree, corroborating the deletion identified by RNA-seq (Fig 2D). Collectively, these results indicate that the entire tree is infected with ANRSV-HK23, whose genome lacks the HCPro1-HCPro2 coding sequence.

#### Determination and analysis of complete genomic sequences of ANSSV-BT23 and ANRSV-HK23

We obtained the full-length genomic sequences of ANSSV-BT23 and ANRSV-HK23 by conventional cloning and sequencing (Figs 3A and 4A). The resulting sequences have been deposited in GenBank database under accession numbers PQ867793 and PQ867792. A pairwise alignment of the genomic sequences from ANSSV-BT23 and ANSSV-BT17 revealed a striking difference: the nucleotide sequence encoding from aspartic acid at position 56 (D56) in HCPro1 to serine at position 578 (S578) in P3 is absent in ANSSV-BT23 (Fig 3B). Notably, the missing region in ANSSV-BT17 is flanked by two short repeated sequences (‘GACGA’), suggesting that the deletion of nearly the entire HCPro1-HCPro2 may have resulted from a recombination event during viral replication. Apart from this major deletion, a total of 41 nucleotide differences were identified between the two sequences, with 14 resulting in missense mutations leading to amino acid changes (four in P3, three in NIa-Pro, NIb and CP, and one in CI) (Fig 3C). Whether these amino acid changes affect viral fitness remains to be determined.

**Fig 3.**
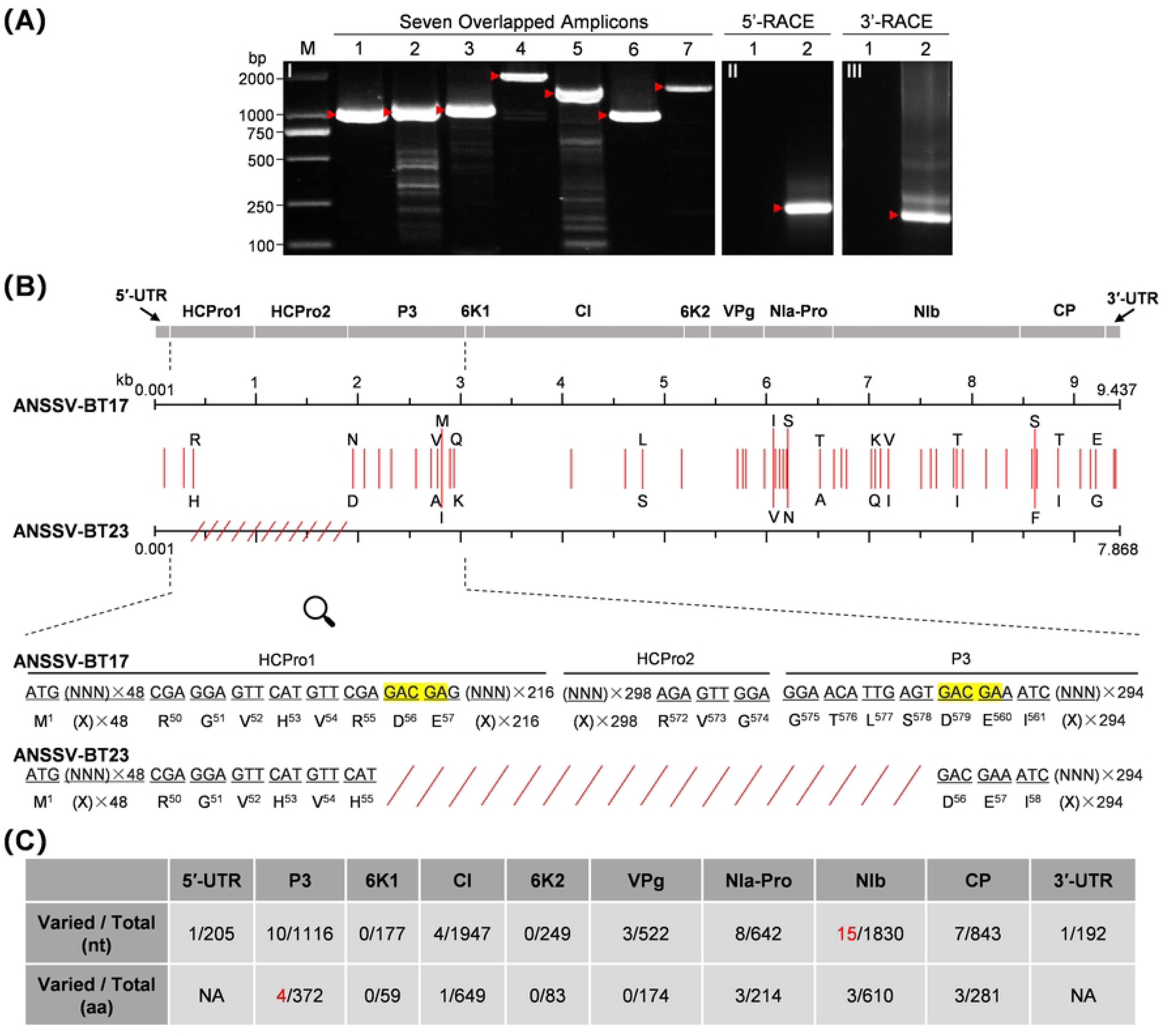
Cloning, sequencing and analysis of the complete genome of ANSSV-BT23. (A) Overlapped amplicons covering the entire genome of ANSSV-BT23. Panel I, RT-PCR amplification of seven overlapping fragments (lanes 1-7) spanning nearly the full-length genome of ANSSV-BT23. Panels II and III, amplicons deriving from 5’-RACE and 3’-RACE, respectively. Lanes 1 and 2 in panels II and III correspond to RNA extracted from healthy and ANSSV-BT23-infected areca palm trees, respectively. (B) Sequence comparison between the genomes of ANSSV-BT17 and ANSSV-BT23. The mutated nucleotides are indicated by vertical red lines. The amino acid changes resulting from missense mutations are shown. The missing region in ANSSV-BT23 is represented by red slant lines. The sequence alignment of HCPro1-to-P3 region highlights the missing sequence in ANSSV-BT23, with the duplicated 5-nt sequences flanking the deletion shaded in yellow. (C) Summary of nucleotide and amino acid variations in each viral cistron when comparing ANSSV-BT17 and ANSSV-BT23.

**Fig 4.**
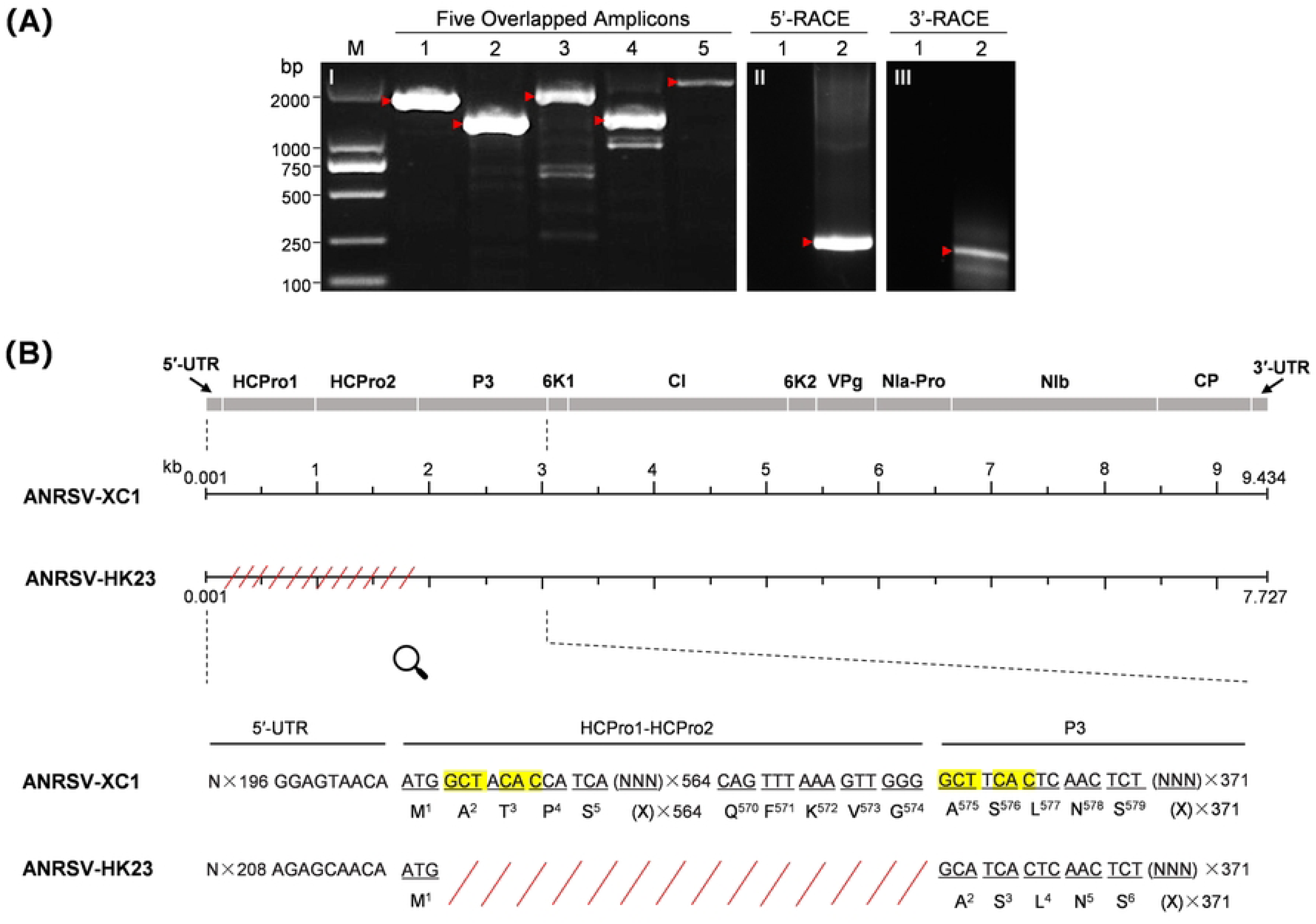
Cloning, sequencing and analysis of the complete genome of ANRSV-HK23. (A) Overlapped amplicons covering the entire genome of ANSSV-BT23. Panel I, RT-PCR amplification of five overlapping fragments (lanes 1-5) spanning nearly the full-length genome of ANRSV-HK23. Panels II and III, amplicons deriving from 5’-RACE and 3’-RACE, respectively. Lanes 1 and 2 in panels II and III correspond to RNA extracted from healthy and ANRSV-HK23-infected areca palm trees, respectively. (B) Sequence comparison between the genomes of ANRSV-HK23 and ANRSV-XC1. The mutated nucleotides are indicated by vertical red lines. The amino acid changes resulting from missense mutations are shown. The missing region in ANRSV-HK23 is represented by red slant lines. The sequence alignment of 5**′**-UTR-to-P3 region highlights the missing sequence in ANRSV-HK23, with the duplicated 6-nt sequences flanking the deletion shaded in yellow. (C) Summary of nucleotide and amino acid variations in each viral cistron when comparing ANSSV-BT17 and ANSSV-BT23.

Blast analysis revealed that the genomic sequence of ANRSV-HK23 shares the highest identity with that of ANRSV-XC1 (MH395371), showing 91.2% nt identity at the whole-genome level. Therefore, the genomic sequence of ANRSV-XC1 was retrieved for the pairwise alignment. The results showed that the complete coding sequence of HCPro1-HCPro2, with the exception of initiation codon, is absent in ANRSV-HK23 (Fig 4B). Two highly-similar short sequences (‘GCTACAC’ vs ‘GCTTCAC’) flank the HCPro1-HCPro2 coding region in ANRSV-XC1 (Fig 4B). We speculate that the parental isolate of ANRSV-HK23 originally had two identical short sequences that underwent a recombination event. Unfortunately, leaf samples were not collected from that particular tree in 2021, preventing us from testing this hypothesis. Based on results described so far, we conclude that ANSSV-BT23 and ANRSV-HK23 have lost the HCPro1-HCPro2 coding sequences from their genome, which correlates with a strong attenuation of viral symptoms in areca palm. The presented data also support the idea that these genomic deletions result from recombination events during viral replication.

#### The absence of HCPro1-HCPro2 attenuates viral infection in *N. benthamiana*

As shown above, ANSSV-BT17, which expresses HCPro1-HCPro2, accumulates at high levels in areca palm, resulting in severe infection symptoms. Conversely, leader protease-less ANSSV-BT23 and ANRSV-HK23 accumulate at much lower levels and produce mild or no symptoms. This phenomenon led us to hypothesize that the absence of HCPro1-HCPro2 attenuates viral infection, while still allowing viral replication and movement. To test this idea, we developed full-length cDNA clones of ANSSV-BT23 and ANRSV-HK23 (pSS-BT23 and pRS-HK23, respectively), in which the GFP coding sequence was inserted between that of NIb and CP to facilitate the observation of virus accumulation under UV light (Figs 5A, S1 and S2). In addition, we developed two viral clones, pSS-BT23-HCPro1-2^17^ and pRS-HK23-HCPro1-2^ZYZ^, for which the absent regions in ANSSV-BT23 and ANRSV-HK23 were restored with the corresponding sequences from ANSSV-BT17 and ANRSV-ZYZ (MZ209276), respectively (Fig 5A).

**Fig 5.**
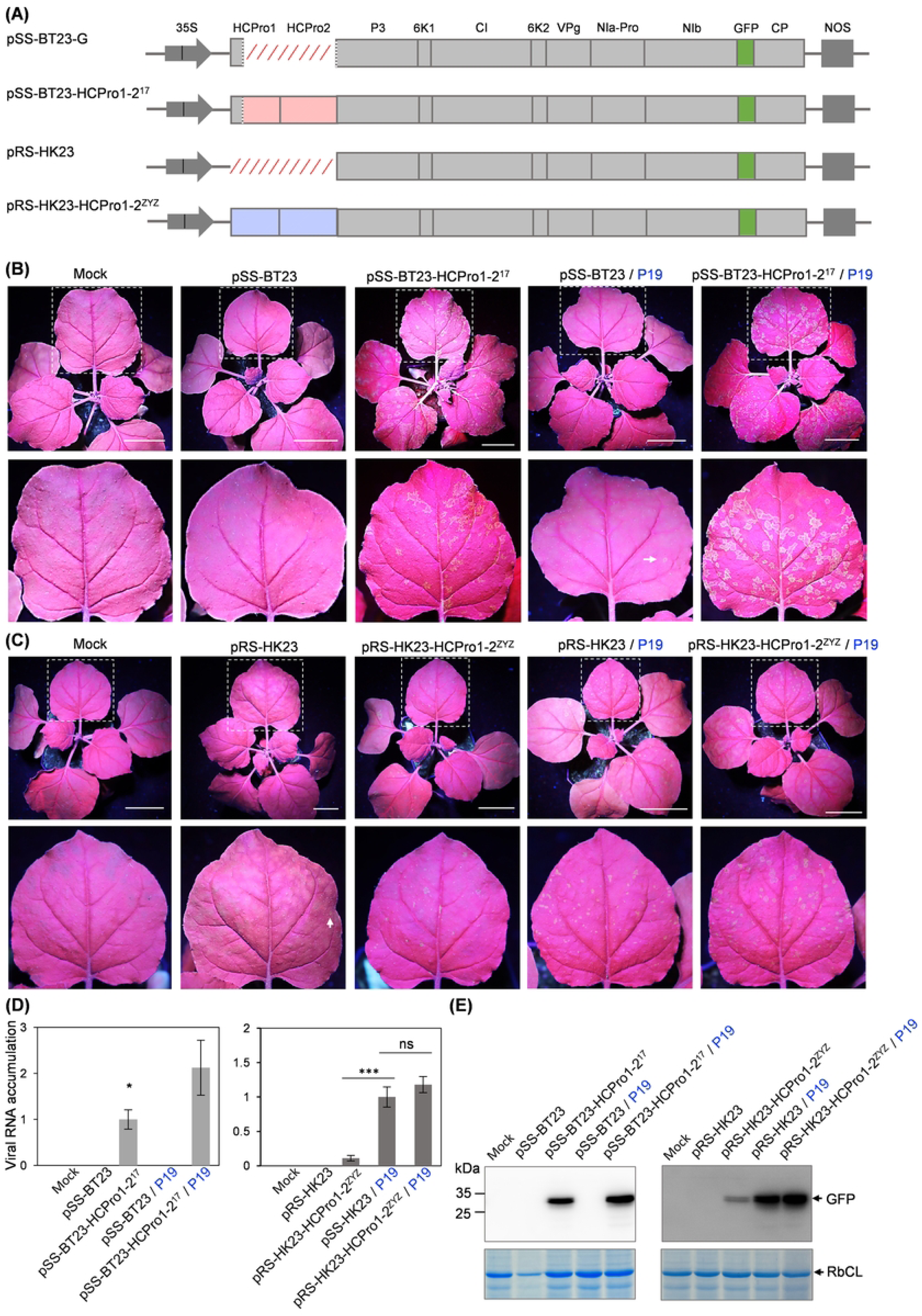
The absence of HCPro1-HCPro2 in ANSSV-BT23 and ANRSV-HK23 attenuates viral infection. **(A)** Schematic diagrams of the indicated virus clones. The nearly complete (for pSS-BT23) and complete (for pRS-HK23) deletion of HCPro1-HCPro2 is indicated by red slant lines. For the hybrid clones, the regions indicated by red and blue rectangles are from ANSSV-BT17 and ANRSV-ZYZ, respectively. **(B)** Inoculation of pSS-BT23 and pSS-BT23-HCPro1-2^17^ in *N. benthamiana*. The indicated clones were agroinoculated into *N. benthamiana* plants (OD_600_ = 0.5 / clone). Alternatively, the indicated plasmids were agoinoculated along a plasmid expressing TBSV P19 (final OD_600_ = 0.5 / clone) in the infiltrated leaf. Representative plants were photographed at 18 days post-inoculation (dpi) under a UV lamp. Leaves indicated with rectangles in white dotted lines are enlarged. White arrows in (B) and (C) indicate scattered fluorescence spots. Bars, 5 cm. **(C)** Inoculation of pRS-HK23 and pRS-HK23-HCPro1-2^ZYZ^. Idem to (B), except that representative plants were photographed at 20 dpi under a UV lamp. **(D)** Real-time RT-qPCR analysis of accumulation levels corresponding to the indicated viruses at 18 dpi (left) or 20 dpi (right). Two primer sets, SS23-qPCR-F/SS23-qPCR-R and RS23-qPCR-F/RS23-qPCR-R (S1 Table) targeting viral *CPs*, were designed to quantify the accumulation of ANSSV-BT23 / ANSSV-BT23-HCPro1-2^17^, and ANRSV-HK23 / ANRSV-HK23-HCPro1-2^ZYZ^, respectively. A newly expanded leaf per plant was sampled for the assay. Error bars represent standard errors from three biological replicates. Statistically significant differences were determined by an unpaired two-tailed Student’s *t* test: ***, *P*<0.001; *, 0.01<*P*<0.05; ns, *P*>0.05. **(E)** Immunoblot detection of GFP at 18 dpi (left) and 20 dpi (right) in protein samples from plants infected with the indicated viruses. Coomassie blue staining of RbCL was used as a loading control.

The infectivity of pSS-BT23 and pSS-BT23-HCPro1-2^17^ was assessed in *Nicotiana benthamiana* plants, as multiple attempts to inoculate arepavirus clones in areca palm have been unsuccessful. The viral clones were agroinoculated into plants either alone or together with an agrobacterium strain that harbours a plasmid expressing tomato bushy stunt virus (TBSV) P19 (Li et al., 2014), a potent RSS that ensures viral protein expression at least in the inoculated leaves. In two independent experiments, all 24 plants agroinoculated with either pSS-BT23-HCPro1-2^17^ (10 plants) or pSS-BT23-HCPro1-2^17^ with P19 (14 plants) exhibited strong green fluorescence in upper non-inoculated leaves, indicating systemic viral infection (Fig 5B). This observation was further confirmed by real-time RT-qPCR and immunoblot analysis (Fig 5D and 5E). In contrast, only four out of 13 plants co-inoculated with pSS-BT23 and P19 displayed scattered fluorescence spots (Fig 5B). Among the remaining plants inoculated with either pSS-BT23 and P19 (9 plants), or pSS-BT23 alone (18 plants), no GFP signal was observed in upper non-inoculated leaves (Fig 5B). Real-time RT-qPCR and immunoblot analysis confirmed the absence of viral infection in these plants (Fig 5D and 5E).

Subsequently, we compared the infectivity of pRS-HK23 and pRS-HK23-HCPro1-2^ZYZ^. In two independent experiments, clear fluorescence spots were observed in all 26 plants either inoculated with pRS-HK23-HCPro1-2^ZYZ^ (12 plants) or co-inoculated with pRS-HK23-HCPro1-2^ZYZ^ and P19 (14 plants) (Fig 5C), indicating that ANRSV-HK23-HCPro1-2^ZYZ^ can systemically infect *N. benthamiana* regardless of the presence of P19 in the inoculated leaves (Fig 5C). This observation was further confirmed by real-time RT-qPCR and immunoblot analysis (Fig 5D and 5E). Similarly, all 13 plants co-inoculated with pRS-HK23 and P19 also exhibited systemic infection, as indicated by GFP fluorescence, real-time RT-qPCR and immunoblot analysis (Fig 5C, 5D and 5E). However, only five out of 31 plants inoculated with pRS-HK23 alone displayed scattered fluorescence spots (Fig 5C).

Together, these results indicate that the absence of HCPro1-HCPro2 in ANSSV and ANSRV significantly attenuates viral infectivity, and that the *trans*-supplementation with a strong RSS in inoculated leaves partially compensates for this defect. A key conclusion from these data is that the lack of HCPro1-HCPro2 does not completely abolish viral replication, nor does it prevent both cell-to-cell and long-distance movement.

#### The loss of HCPro1-HCPro2 weakens viral-mediated RNA silencing suppression and other essential functions

Since *trans*-supplementation with P19 in inoculated leaves alleviated viral defects, we hypothesized that the attenuation of arepaviruses lacking HCPro1-HCPro2 might, at least in part, result from a defect in RNA silencing suppression. Indeed, we have previously demonstrated that HCPro2, either alone or as part of the HCPro1-HCPro2 tandem, functions as an RSS in ANSSV and ANSRV (Qin et al., 2021, 2024). To test the role of RNA silencing in the attenuation of viruses without HCPro1-HCPro2, we used a transgenic *N. benthamiana* line, here referred to as *dcl2/4*, in which the expression of Dicer-Like Protein 2 (DCL2) and 4 (DCL4) has been strongly reduced by RNAi (Dadami et al., 2013). In two independent experiments, pSS-BT23 and pSS-BT23-HCPro1-2^17^ were inoculated into 16 and 11 *dcl2/4* plants, respectively. As a control, pSS-BT23 and pSS-BT23-HCPro1-2^17^ were also inoculated into ten wild-type *N. benthamiana* plants per construct.

Consistent with the results presented in the previous section, the detection of GFP signal in upper non-inoculated leaves indicated that ANSSV-BT23-HCPro1-HCPro2^17^, but not ANSSV-BT23, successfully infected wild-type *N. benthamiana* plants (Fig 6A). Notably, ANSSV-BT23 infection was restored in all inoculated *dcl2/4* plants (Fig 6A). These findings were further corroborated by real-time RT-qPCR and immunoblot analyses (Fig 6C and 6D). However, when compared viral accumulation based on GFP signal, real-time RT-qPCR, and immunoblot analysis between ANSSV-BT23 plants and ANSSV-BT23-HCPro1-2^17^ in *dcl2/4* plants, the latter exhibited higher levels (Fig 6B, 6C and 6D). This result indicates that ANSSV-BT23 does not reach the same fitness level as ANSSV-BT23-HCPro1-2^17^ in plants deficient in RNA silencing. Therefore, it is likely that the HCPro1-HCPro2 tandem plays additional roles beyond RNA silencing suppression.

**Fig 6.**
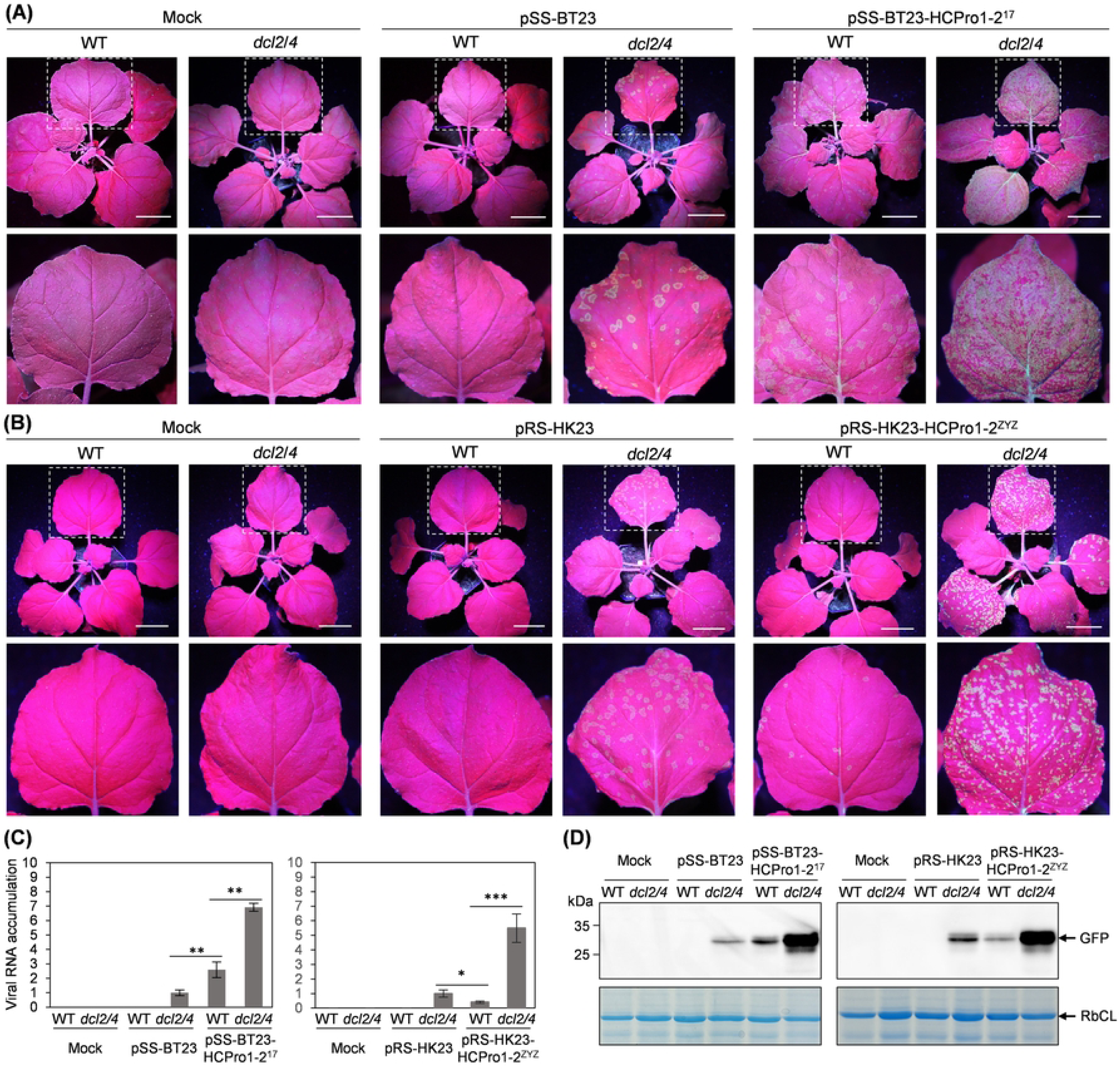
The absence of HCPro1-HCPro2 compromises RNA silencing suppression activity and affects other critical functions. **(A)** Inoculation of pSS-BT23 or pSS-BT23-HCPro1-2^17^ in wild-type (WT) and *dcl2/4 N. benthamiana* plants. The indicated clones were agroinoculated into plants (OD_600_ = 0.5 / clone). Representative plants were photographed at 20 days post-inoculation (dpi) under a UV lamp. Leaves indicated with rectangles in white dotted lines are enlarged. Bars, 5 cm. **(B)** Inoculation of pRS-HK23 or pRS-HK23-HCPro1-2^ZYZ^. Idem to (B). **(C)** Real-time RT-qPCR analysis of viral genomic RNA accumulation corresponding to the indicated viruses at 20 dpi. Two primer sets, SS23-qPCR-F/SS23-qPCR-R and RS23-qPCR-F/RS23-qPCR-R (S1 Table), were used to quantify the accumulation of ANSSV-BT23 / ANSSV-BT23-HCPro1-2^17^, and ANRSV-HK23 / ANRSV-HK23-HCPro1-2^ZYZ^, respectively. A newly expanded leaf per plant was sampled for the assay. Error bars represent standard errors from three biological replicates. Statistically significant differences were determined by an unpaired two-tailed Student’s *t* test: ***, *P*<0.001; **, 0.01<*P*<0.001; *, 0.01<*P*<0.05. **(D)** Immunoblot detection of GFP abundance at 20 dpi in protein samples from plants infected with the indicated viruses. Coomassie blue staining of RbCL was used as a loading control.

The infectivity of both ANRSV-HK23 and ANRSV-HK23-HCPro1-2^ZYZ^ was also examined in wild-type and *dcl2/4 N. benthamiana* plants. Briefly, the results of this experiment were similar to those obtained with ANSSV-BT23 and ANSSV-BT23-HCPro1-HCPro2^17^ (Fig 6A, 6C and D), as the infectivity of the attenuated ANRSV-HK23 is restored in all the inoculated *dcl2/4* plants. As observed with ANSSV derivatives, when comparing the fitness of ANRSV-HK23 and ANRSV-HK23-HCPro1-2^ZYZ^ in *dcl2/4* plants, we found that the accumulation of the former was much lower than that of the latter (Fig 6B-6D). This result further supports the idea that the HCPro1-HCPro2 tandem plays additional roles beyond RNA silencing suppression. In fact, our recent report on the role of HCPro2 in virus cell-to-cell movement provides some insights into this matter (Qin et al., 2024).

#### A field survey of trees revealed a potential intermediate stage in the evolution of ANRSV, where full-length isolates coexist with leader protease-less variants

The results described above indicate that two areca palm trees are independently infected with arepaviruses carrying genomic deletions in the HCPro1-HCPro2 coding sequence. To determine the prevalence of this phenomenon in other naturally growing trees, we conducted a field survey. A total of 52 adult areca palm trees growing in Danzhou, Hainan were selected for the study. First, plants were classified into five groups based on symptom severity (S3 Fig). Leaf samples from all these trees were independently collected and used for total RNA extraction, followed by RT-PCR analysis. For this purpose, a pair of degenerate primers, 5UTR-DP-F1/P3-DP-R1, was designed to flank the HCPro1-HCPro2 coding region from ANSSV and ANRSV (S1 Table and S4 Fig). Additionally, another pair, CP-DP-F2/CP-DP-R1, was designed to target the CP coding sequence from ANSSV and ANRSV (S5 Fig). These primers were used for subsequent assays.

An amplicon of 325 bp was produced using the primer pair targeting the CP coding region in RNA samples from all symptomatic trees, but not in those from asymptomatic ones (S3 Fig). Sanger sequencing confirmed that all symptomatic trees were infected with ANRSV. Similarly, when using the primer pair flanking the HCPro1-HCPro2 coding sequence, all samples from symptomatic trees yielded a long amplicon of approximately 2100 bp (S5 Fig), indicating the presence of full-length ANRSV isolates. Importantly, the use of the primer pair flanking the HCPro1-HCPro2 coding sequence also generated a short amplicon in RNA samples from infected trees, particularly those with severe foliage necrosis (i.e., #8, #10, #15, #23), including the one infected with ANRSV-ZYZ (S3 Fig). Short amplicons from each RT-PCR reaction were independently cloned, and three independent colonies per plate were used for plasmid preparation and sequencing. This analysis demonstrated that these amplicons corresponded to highly similar fragments of different sizes, all lacking the HCPro1-HCPro2 coding sequence. Moreover, pairwise alignment of these short amplicons revealed that (i) the deleted region either spans from the 5’ end of the P1 coding sequence (in most cases) or from the final sequence of the 5’ UTR to the coding sequence around the HCPro2/P3 junction (Figs 7 and S6); (ii) all deleted regions are flanked by two short repeated sequences in the corresponding parental virus, supporting the idea that truncated variants originate through recombination; (iii) half of the isolates carrying the HCPro1-HCPro2 deletion maintain the translational frame (Figs 7 and S6), as in ANSSV-BT23 and ANSSV-HK23.

**Fig 7.**
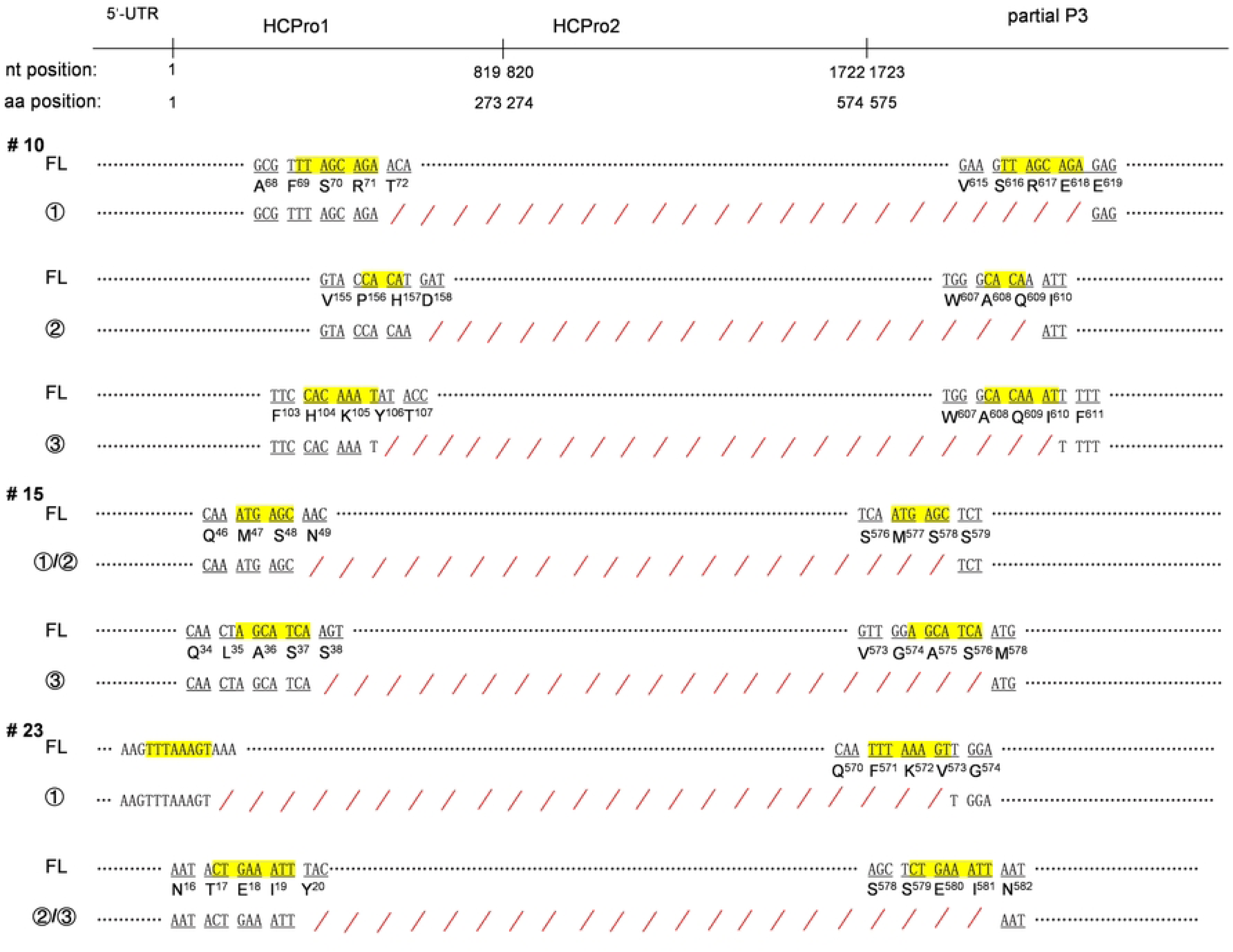
Pair-wise alignments of full-length and truncated ANRSV isolates found in the same tree. For trees #10, #15 and #23, both the long and short amplicons (S3 Fig) were cloned. Plasmids from three independent colonies per plate were sequenced. Sequence from clones corresponding to the long amplicons were identical across samples. Nucleotide sequences of the long (FL) and various short (rounded numbers) amplicons were aligned. Short repeated sequences flanking the deleted fragments in the long amplicon are highlighted in yellow, while deleted regions in the short amplicons are marked with red slant lines.

Taken together, these results suggest that some of these trees, particularly those suffering from severe infection, are in a transitional stage of plant-virus coevolution, in which HCPro1-HCPro2 deleted variants have just begun to emerge. Based on all the results presented in this study, it would be expected that, in the next stages, only the full-length virus disappears from these trees. If this is the case, then we should identify plausible pathways explaining how the deleted variants are able to outcompete the full-length virus.

## Discussion

Recombination events and genomic mutations are common phenomena in viruses, driving their evolution and diversification (Sztuba-Solińska et al., 2011; Pérez-Losada et al., 2015; Stern and Andino., 2016). To the best of our knowledge, this study presents the first report of potyvirids losing a significant portion of their genome, likely via recombination, resulting in self-attenuation and long-time persistence. In the specific cases described here, viruses lacking leader proteases are produced. Interestingly, shorter viral RNA genomes, derived from full-length arepaviruses, resemble the RNA1 segment from members of the *Bymovirus* genus, which possess bipartite genomes consisting of RNA1 and RNA2. This similarity raises the intriguing possibility that arepaviruses might have evolved by fragmenting their originally monopartite genomes into two RNA segments. However, our attempts to detect RNA2-like genomes, expected to contain HCPro1-HCPro2 coding sequence (or similar derivatives), in our RNA-seq data were unsuccessful (not even a single read showing similarity to the HCPro1-HCPro2 coding sequences was detected). This result indicates that, in the long term, the viral entities infecting areca palm trees remain monopartite and lack the HCPro1-HCPro2 coding sequence.

Our data support the idea that genomic deletions in arepaviruses arise through recombination during the replication of full-length parental viruses (Figs 7 and S6). Remarkably, recombination is the mechanism responsible for the generation of D-RNAs from their parental viruses. The prolonged passaging of viruses at a high multiplicity of infection within a single host is a key factor in the generation of D-RNAs (Budzyńska et al., 2022; Joanna Sztuba-Solinska et al., 2011; Simon et al., 2004). Consistently, arepaviruses with genomic deletions appear to be produced in trees exhibiting severe foliage necrosis and accumulating high level of the parental virus (Figs 1 and S3). We therefore propose that the generation of arepavirus variants harboring genomic deletions follows similar principles to those underlying D-RNA formation. However, despite the here-mentioned similarities, D-RNAs are small in size and lack the coding sequences for essential factors involved in replication, encapsidation and/or movement. As a result, they strictly depend on the presence of full-length parental viruses for persistence. This key distinction leads us to conclude that arepaviruses harboring large genomic deletions are not D-RNAs.

Two types of leader proteases, P1 and HCPro, are produced by all monopartite potyvirids, although their arrangement within the polyprotein is variable. Several evolutionary events, in particular recombination and gene duplication, have been implicated in shaping the hypervariable 5’-proximal regions of monopartite potyvirid genomes (Valli et al., 2007; Qin et al., 2021; Rodamilans et al., 2021; Pasin et al., 2022; Gibbs et al., 2020). These two proteins are believed to be among the most divergent potyvirid factors, suggesting that leader proteases play a crucial role in host range expansion and adaption (Valli et al., 2007; Shan et al., 2015; Carbonell et al., 2012). In member of the *Bymovirus* genus, RNA2 resembles the 5’-terminal region of monopartite potyvirids, encoding two mature proteins: P1 (a homolog of HCPro) and P2, both of which act as RSSs, at least in the case of wheat yellow mosaic virus (Chen et al., 2023, 2024). Additionally, P2 is directly involved in vector transmission (Adams et al., 2001). Remarkably, during successive passages by mechanical inoculation, RNA2 in bymoviruses also loses a genomic fragment. In these cases, the deleted fragment encodes a P2 domain, yet its deletion does not lead to viral attenuation, but abolishes transmission (Dessens et al., 1995; Schenk et al., 1995; Timpe & Kühne, 1995; Zheng et al., 2002). The contrasting effects of genomic fragment loss in bymoviruses versus arepaviruses can be explained by the fact that bymoviruses carrying the deletion still express the full-length P1 protein. This would prevent the action of the host antiviral mechanism based on RNA silencing, ensuring that bymoviruses, unlike arepaviruses, remain fully competent.

Arepaviruses with deletions in their genomes are significantly attenuated, as evidenced by (i) the low number of reads in RNA-seq data from infected areca palm trees (Figs 1 and 2) and (ii) virus accumulation assessments in *N. benthamiana* plants (Figs 5 and 6). Our findings on arepavirus self-attenuation led us to propose a comprehensive scenario explaining how and why these variants emerge and, intriguingly, persist in infected trees while the fully competent parental variants are eliminated (Fig. 8): (a) arepaviruses are transmitted by an as-yet-unknown natural vector into healthy trees, initiating the infection; (b) viruses spread to new leaves, triggering disease symptoms alongside the activation of diverse antiviral mechanisms; (c) one of these mechanisms is activated by, or targets, the HCPro proteins, as supported by previous findings in other potyvirids (Nakahara et al., 2012; Hafrén et al., 2018; Moury et al., 2011); (d) while full-length viruses are eliminated from the infected tree, attenuated viral variants lacking *HCPro1-HCPro2* emerge through recombination; (e) both the full-length and attenuated viruses coexist in the same tree (supported by results shown in Figs 7 and S6); (f) eventually, only the virus with the genomic deletion, which remains asymptomatic, persists in the tree. Given the key role of HCPro during vector transmission in potyvirids (Valli et al., 2018), it is likely that arepaviruses lacking HCPro1-HCPro2 are not transmitted between trees. There are at least two non-mutually exclusive possibilities explaining how self-attenuated arepaviruses lacking the RSS HCPro2 persist in trees for long time: (i) VPg in arepaviruses might also possess some RNA silencing suppression activity, as reported for certain potyviruses (Rajamaki & Valkonen, 2009; Cheng & Wang, 2017), allowing viruses without HCPro2 to persist; and (ii) a potent RSS may be strictly required at the onset of infection, but not in the long term. This idea is supported by findings in Fig 5, where HCPro2-deficient viruses were able to spread over long distances in some plants when complemented with P19 in inoculated leaves. In any case, the intriguing possibility that trees might not actively eliminate self-attenuated arepaviruses for a specific reason might provide an evolutionary context for their persistence. In fact, we propose that the presence of self-attenuated arepaviruses is beneficial to trees by conferring cross-protection (Ziebell & Carr, 2010; Zhang & Qu, 2016) against new infections from pathogenic parental strains or closely related viruses, such as ANRSV2, a newly-proposed species within the *Arepavirus* genus (Pandian et al., 2024). Indeed, a recent report has revealed that a mycovirus is co-opted by plants to mediate broad-spectrum fungal resistance (Wang et al., 2024).

**Fig 8.**
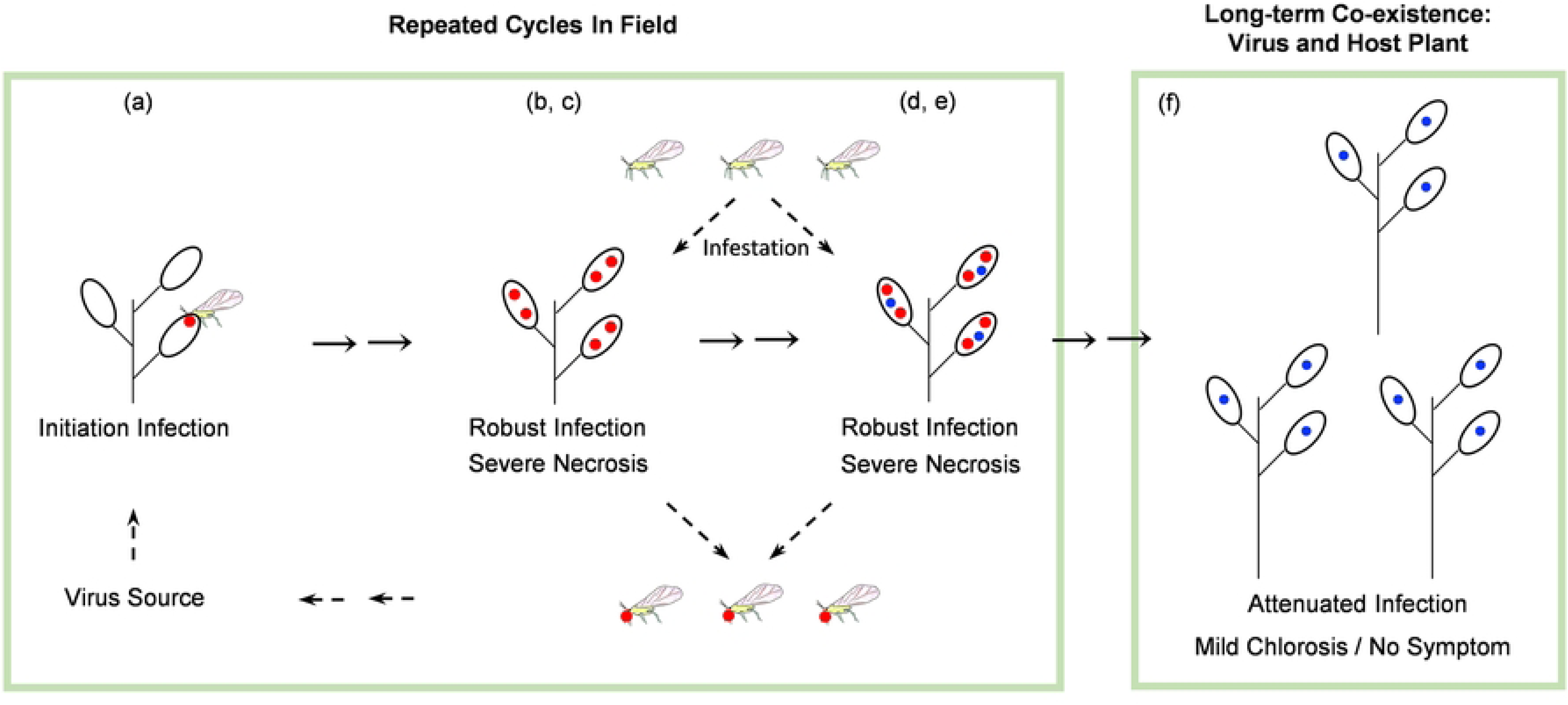
A model illustrating the evolution of arepaviruses in infected trees. The full-length parental viruses and isolates lacking HCPro1-HCPro2 are marked with red and blue dots, respectively. See the Discussion section for a detailed description of the evolutionary pathway.

Overall, our findings provide valuable insights into the relationship between viral evolution and the long-term stability of virus-host interactions. Further studies will be necessary to deeply understand virus self-attenuation through recombination, as well as to validate the putative transition from pathogenic to mutually beneficial interaction in areca palm trees infected with arepaviruses.

## Materials and methods

### Plants and virus isolates

The areca palm tree infected by ANSSV-HNBT (Yang et al., 2018) is growing in the field under natural conditions. The virus isolates identified in that tree in 2017 and 2023 are designated here as ANSSV-BT17 and ANSSV-BT23, respectively. A tree infected by ANRSV in Haikou, Hainan, showed severe foliage necrosis in 2021, but it was asymptomatic in 2023. The virus isolates in this tree in 2021 and 2023 are named ANRSV-HK21 and ANRSV-HK23, respectively. Leaf samples from both trees were collected at the indicated time points for RNA-seq, virus genome cloning, and generation of full-length cDNA clones. For the field survey, a total of 52 adult areca palm trees (most of them showing typical foliage necrosis), growing in a small field in Danzhou, Hainan (19.510688°N, 109.488776°E) in August, 2024, together with a previously-stored leaf sample infected with ANRSV-ZYZ (Wang et al., 2021), were selected for further analyses. Seedlings of wild-type and *dcl2/4* (Dadami et al., 2013) *N. benthamiana* plants were maintained in a growth cabinet (16 h-light, 25°C and 8 h-darkness, 23°C, 70% relative humidity).

### Plasmids

The full-length cDNA clones of pSS-BT23 and pRS-HK23 with a GFP-tag at the NIb-CP junction were developed. For the generation of pSS-BT23, the following procedure was performed (S1 Fig). Step 1: amplification of six fragments with approximately 30-nt overlapping sequence with both flanking fragments. Fragments 1, 3, 4 and 6, covering the entire genome of ANSSV-BT23, were amplified from cDNAs (prepared from ANSSV-BT23-infected areca palm leaves) with primer sets ANSSV-All-F/Intron-SOE-2, Intron-SOE-3/SS-5900R, SS-4200F/SOE-G-2 and SOE-G-5/ANSSV-All-R (S1 Table). The pSS-I-G (Qin et al., 2021) was used as the template to amplify fragments 2 and 5 with the primer sets Intron-SOE-1/Intron-SOE-4 and SOE-G-3/SOE-G-4. Step 2: overlapping PCRs to yield fragments 1-to-3 and 4-to-6. The amplicons of fragments 1-3 or 4-6 were mixed as the template for overlapping PCR with the primer set ANSSV-All-F/SS-5900R (for fragment 1-to-3) or SS-4200F/ANSSV-All-R (fragment 4-to-6). Step 3: the fragment 7, as the backbone, was amplified from pCB301-35S-Nos (Cui & Wang, 2016) with the primer set e35S-R/Nos-F (S1 Table). Step 4: homologous recombination of the fragments 1-to-3, 4-to-6 and 7 to yield pSS-BT23. For the release of GFP from the viral polyprotein, two short nt sequences (GCAAACAAGGAATTCCAA/ATGGA<u>C</u> and GC<u>C</u>AA<u>T</u>AA<u>A</u>GA<u>G</u>TT<u>T</u>CA<u>G</u>/ATGGAT) coding for the same peptide “ANKEFQ/MD” at the NIb-CP junction of ANSSV-BT23 (proteolytically processed by NIa-Pro) were engineered into NIb-GFP and GFP-CP junctions, respectively. Seven nucleotide difference (underlined) between the two sequences were introduced, which make them less likely to undergo recombination during replication.

The pRS-HK23 was generated by following the here indicated steps (S2 Fig). Step 1: amplification of four fragments with approximately 30-nt overlapping sequence with both flanking fragments. Fragments 1-3, covering the entire genome of ANRSV-HK23, are amplified from the cDNAs (prepared from ANRSV-HK23-infected areca palm leaves) with primers sets ANRSV23-All-F/RSV-5466R, RSV-3689F/RS23-SOE-G-1R, and RS23-SOE-G-2F/RS23-All-R (S1 Table). The pRS-G (Wang et al., 2021) was used as the template to amplify fragment 4 with the primer set RS23-SOE-G-1F/RS23-SOE-G-4. Step 2: overlapping PCRs to yield fragments 1-to-2 and 3-to-4. Amplicons of fragments 1 and 2 or 3 and 4 were mixed as the template for overlapping PCR with the primer set ANRSV23-All-F/RS23-SOE-G-1R or RS23-SOE-G-1F/RS23-All-R. Step 3: the fragment 5, as the backbone, was amplified from pCB301-35S-Nos (Cui & Wang, 2016) with the primer set RS23-Nos-F/e35S-R (S1 Table). Step 4: homologous recombination of the fragments 1-to-2, 3-to-4 and 5 to yield pRS-HK23. For the release of GFP from the viral polyprotein, two short nt sequences (GCAAGCAAGGAATTTCAA/ATGGA<u>C</u> and GC<u>G</u>AGCAA<u>A</u>GA<u>G</u>TT<u>C</u>CA<u>G</u>/ATGGAT) coding for the same peptide ‘ASKEFQ/MD’ at NIb-CP junction of ANRSV-HK23 (proteolytically processed by NIa-Pro) were engineered into NIb-GFP and GFP-CP junctions, respectively. Six nucleotide difference (underlined) between the two sequences were introduced, which make them less likely to undergo recombination during viral replication.

Two hybrid virus clones, pSS-BT23-HCPro1-2^17^ and pRS-HK23-HCPro1-2^ZYZ^, were constructed by standard DNA manipulation techniques. For them, the deleted regions in ANSSV-BT23 and ANRSV-HK23 were reintroduced by using the corresponding regions from ANSSV-BT17 and ANRSV-ZYZ. To construct pSS-BT23-HCPro1-2^17^, the region covering nearly the entire HCPro1-HCPro2 in pSS-I-G (Qin et al., 2021) was amplified with the primer set SS23-SOE-HP1-F/SS23-SOE-HP2-P3-R, and its flanking regions in pSS-BT23 were amplified with primer sets pCB301-F/SS23-SOE-HP1-R and SS23-SOE-HP2-P3-F/SS23-1300R (S1 Table). The obtained products were mixed as the template for overlapping PCR with the primer set pCB301-F/SS24-1300R. The resulting fragment was inserted into pSS-BT23 by using *Pme* I/*Sap* I unique sites to generate pSS-BT23-HCPro1-2^17^. To develop pRS-HK23-HCPro1-2^ZYZ^, the entire region of HCPro1-HCPro2 in pRS-G (Wang et al., 2021) was amplified with the primer set RS23-SOE-5UTR-HP1-F/RS23-SOE-HP2-P3-R, and its flanking regions in pRS-HK23 were amplified with primer sets pCB301-F/RS23-SOE-5UTR-HP1-R and RS23-SOE-HP2-P3-F/RSV-5RACE-1R (S1 Table). The obtained products were mixed as the template for overlapping PCR with the primer set pCB301-F/RSV-5RACE-1R. The resulting fragment was inserted into pRS-HK23 by using *Pme* I/*Apa* I unique sites to generate pRS-HK23-HCPro1-2^ZYZ^.

#### Total RNA extraction, RT-PCR, RT-qPCR

Total RNA of newly-developed leaves from symptomatic areca palm trees was prepared with RNAprep Pure Plant Kit (TIANGEN) with a further treatment with DNase I (Thermo Fisher Scientific). First-strand cDNAs were produced with RevertAid First Strand cDNA synthesis kit (Thermo Fisher Scientific) by using random hexamer primers. PCRs were carried out with KOD One PCR Master Mix-BLUE (TOYOBO) by using the indicated primer sets (Supplemental Table S1). For real-time qPCRs, the gene-specific primers were designed by using Primer3Plus (https://www.primer3plus.com/index.html). The qPCRs were conducted with SuperReal Premix Plus (TIANGEN) in the Applied Biosystems QuantStudio 5 machine (Thermo Fisher Scientific).

#### RNA-seq and analysis

Total RNA samples were sent to a company (BIOWEFIND Co., Ltd. in Wuhan, China) for RNA-seq analysis. After rRNA depletion, the NEBNext Ultra Directional RNA Library Prep Kit (NEB) was used to construct libraries, which were further sequenced in a NovaSeq 6000 platform (Illumina). Clean reads were aligned with either ANSSV-BT17 (MH330686) or ANRSV-XC1 (MH395371) genomes by using bowtie2 (Langmead & Salzberg, 2012). The obtained results were visualized by using the Integrative Genomics Viewer (IGV) (Robinson et al., 2011).

#### Cloning, sequencing and annotation of ANSSV-BT23 and ANRSV-HK23

To obtain the complete genomic sequences of ANSSV-BT23 and ANRSV-HK23, seven (for ANSSV-BT23) or five (for ANRSV-HK23) fragments covering nearly the entire genomes were amplified by RT-PCR with the inidicated primer sets (S1 Table). These primers, including the virus-specific primers used in 5’ or 3’ Rapid Amplification of cDNA Ends (RACE), were designed by using RNA-seq data. Both 5’- and 3’-RACEs were performed by using the corresponding kits (Invitrogen). The resulting PCR products were ligated into pTOPO-Blunt vectors (Aidlab Biotechnologies) and three independent positive clones per fragment were sent for Sanger sequencing (Sangon Biotech). The genomes were annotated with reference to the previously-deposited genomic sequences of ANSSV-BT17 (MH330686), ANRSV-XC1 (NH395371) and ANRSV-ZYZ (MZ209276) in GenBank database.

#### Sequence analysis

Nucleotide sequence analyses, including amino acid sequence deduction, ORF identification, sequence assembly and pairwise sequence alignment were performed with the Lasergene software package version 7.1 (GATC Biotech). Multiple sequence alignment was conducted with Clustal Omega (https://www.ebi.ac.uk/jdispatcher/msa/clustalo). The resulting files were visualized via the online tool ESPript 3.0 (Robert & Gouet, 2014).

#### Agroinfiltration

Seedlings of wild-type and the *dcl2/4 N*. *benthamiana* plants at 3- to 5-leaf stage were used for the infectivity test of the indicated virus clones. Viruses were inoculated into newly expanded leaves via agroinfiltration (*Agrobacterium* strain GV3101), following a previously-described protocol (Wang et al., 2021).

### GFP fluorescence observation

Plants inoculated with GFP-tagged viruses were monitored by using a LUYOR-3410 hand-held UV lamp (LUYOR). Representative plants were photographed in a dark room under UV light.

### Immunoblot

Immunoblot assays were performed by following a previously-described protocol (Hu et al., 2023). Anti-GFP polyclonal antibody (Abcam) and a goat anti-rabbit immunoglobulin antibody (Abcam) conjugated to horseradish peroxidase were used as the primary and secondary antibodies, respectively. An enhanced chemiluminescence detection reagent (Thermo Fisher Scientific) was used as the substrate, and signals were observed by using a multifunctional chemiluminescence imager apparatus (JingYi Technology).

## Acknowledgments

We thank Drs. Juan Antonio Garcia (Spanish National Centre for Biotechnology), Guanwei Wu (Ningbo University), and Wenping Qiu (Missouri State University) for critical suggestions. We are also grateful to Kriton Kalantidis for providing seeds from the *dcl2/4* line. This work was supported by grants from the Key Research and Development Project of Hainan Province (ZDYF2025XDNY073 to HC) and National Natural Science Foundation of China (32360651 to ZD). Work in the laboratory of AAV is supported by a grant from MICIU/AEI/10.13039/501100011033/ and FEDER/UE (PID2022-139314OB-I00).

## Supplemental Figure legends

**S1 Table.** Primers used in this study.

**Fig. S1** Generation of the GFP-tagged clone of ANSSV-BT23 (pSS-BT23). Purple and blue rectangles represent a 220-bp intron 2 of *NiR* gene (*Phaseolus vulgaris*) and the complete GFP-coding sequence, respectively. Vertical white lines inside and outside P3-to-CP in the ANSSV-BT23 genome represent NIa-Pro cleavage sites and initiation / stop codons, respectively.

**Fig. S2** Generation of GFP-tagged clone of ANRSV-HK23 (pRS-HK23). Blue rectangles represents the complete GFP-coding sequence. Vertical white lines inside and outside P3-to-CP represent NIa-Pro cleavage sites, and initiation / stop codons, respectively.

**Fig. S3** RT-PCR analysis demonstrates the co-existence of full-length and leader protease-less isolates in the same trees growing in the field. Adult trees in the field were classified into five groups based on disease symptomatology: -, symptomless; CS, chlorotic spots in top leaves; the symbols *, ** and *** indicate that approximately one third, half, and two third of total leaves in the trees display severe necrotic spots, respectively. A newly-expanded leaf from each tree was sampled for total RNA extraction, and RT-PCR with the indicated primer sets was carried out. The presence of ANSSV / ANRSV was assessed with a pair of degenerate primers: CP-DP-F2 / CP-DP-R1 (middle panels). The region spanning HCPro1-HCPro2 was amplified with another primer of degenerated primers: 5UTR-DP-F1 / P3-DP-R1 (upper panels). The detection of *AcActin* transcripts in each sample was used as internal control (lower panels). A leaf from an areca palm tree infected with ANRSV-ZYZ and showing severe foliage necrosis was harvested in Dingan, Hainan, in 2020 (Wang et al., 2021). This tissue was also used to prepare total RNA for RT-PCR.

**Fig. S4** Designing degenerated primers to amplify genomic fragments from 5’UTR to P3 coding sequence. Multiple alignment of 5’ UTR (upper) or partial P3 sequences (lower) of ANSSV and ANRSV isolates, highlighting the regions corresponding to the pair of degenerate primers 5UTR-DP-F1 and P3-DP-R1. Except for the sequences of ANSSV-BT23 (PQ867792) and ANRSV-HK23 (PQ867793) obtained in this study, the remaining ones were retrieved from the GenBank database: ANSSV-HNBT (namely ANSSV-BT17 in this study) (MH330686), ANRSV-XC1 (MH395371), and ANRSV-ZYZ (MZ209276). Identical residues are shown with white letters in red background, whereas conserved substitutions are displayed with red letters in blue boxes.

**Fig. S5** Designing degenerated primers to amplify genomic fragments from the CP coding sequence. Multiple alignment of *CP* sequences from ANSSV and ANRSV isolates, highlighting the regions corresponding to the pair of degenerate primers CP-DP-F2 and CP-DP-R1. Except for the sequences of ANSSV-BT23 and ANRSV-HK23 obtained in this study, the others were retrieved from the GenBank database: ANSSV-BT17 (MH330686), ANRSV-XC1 (MH395371), ANRSV-NY2 (MH395387), ANRSV-MK1 (MH395380), ANRSV-NP2 (MH425893), ANRSV-MK4 (MH395383), ANRSV-MK3 (MH395382), ANRSV-MK2 (MH395381), ANRSV-NY1 (MH395386), ANRSV-NY3 (MH425891), ANRSV-ZYZ (MZ209276), ANRSV-DH1 (MH395375), ANRSV-DH5 (MH395378), ANRSV-DH2 (MH395376), ANRSV-DH4 (MH395377), ANRSV-NH2 (MH395385), ANRSV-NH1 (MH395384), ANRSV-SB6 (MH395393), ANRSV-LG2 (MH395379), ANRSV-XC2 (MH425890), ANRSV-NY4 (MH395388), ANRSV-NY5 (MH395389), ANRSV-SB4 (MH395391), ANRSV-SB5 (MH395392), ANRSV-DA1 (MH395372), ANRSV-DA2 (MH395373), ANRSV-DA3 (MH395374), ANRSV-DAT (MW282956), ANRSV-NP3 (MH425894), ANRSV-SB3 (MH395390). Identical nucleotides are shown with white letters in red background, whereas conserved substitutions are displayed with red letters in blue boxes.

**Fig. S6** Pair-wise alignment of full-length and truncated ANRSV isolates found in the same trees. For tree #8 and the one infected with ANRSV-ZYZ, both the long and short amplicons (S3 Fig) were cloned. Plasmids from three independent colonies per plate were sequenced. Sequence from clones corresponding to the long amplicons were identical across samples. Nucleotide sequences of the long (FL) and various short (rounded numbers) amplicons were aligned. Short repeated sequences flanking the deleted fragments in the long amplicon are highlighted in yellow, while deleted regions in the short amplicons are marked with red slant lines.

## Notes

### Competing Interest Statement

The authors have declared that no competing interests exist.

